# Bacterial ancestry of the mitochondrial ATP exporter

**DOI:** 10.64898/2026.03.31.715626

**Authors:** Jotin Gogoi, Rajan Sankaranarayanan

## Abstract

Mitochondria originated through endosymbiosis of an Alphaproteobacterium within an Asgard archaeal host, with ATP export to the cytosol being a key driver for the organelle integration. The mitochondrial ATP/ADP carrier (AAC), a member of the SLC25 family, performs this critical function, whose evolutionary origin was not known due to the absence of any known prokaryotic homologues and was therefore termed a eukaryotic innovation. Here, using protein tertiary structure search combined with comprehensive sequence analyses, we identify conserved bacterial inner membrane transporters, CysZ and YihY, as putative homologues of mitochondrial AAC. CysZ and YihY are structurally similar to AAC, albeit with a circular permutation of one of the six transmembrane helices. Strikingly, we could identify the conserved MCF motif—a characteristic feature of the SLC25 family—in the bacterial sulfate transporter CysZ, suggesting a common ancestry. Together, our results identify a bacterial origin for the mitochondrial ATP exporter, thus resolving a long-standing question in mitochondrial evolution and a key step required for the emergence of eukaryotic cell complexity.

## Introduction

Eukaryotic cell has evolved from its Asgard archaeal ancestor around 2 billion years ago upon symbiosis of an Alphaproteobacterial endosymbiont that became mitochondria (**Vosseberg et al., 2024**). The energy budget afforded by mitochondria enabled the emergence of the profound eukaryotic cell complexities, which the prokaryotic cells could never evolve in around 4 billion years of evolution (**Lane and Martin, 2010**). The surplus production and supply of ATP to the host cell incentivised the integration of the bacterial endosymbiont leading to its transformation into mitochondria that afforded the energetic cost for evolution of eukaryotic cell complexities **(Roger et al., 2017; Martin, 2025; Speijer, 2025)**. Thus, the evolution of the mitochondrial ATP exporting translocase, ATP/ADP carrier (AAC), is a foundational step in all the models of the emergence of mitochondria and eukaryotic cell (**Gray, 2014**). Notably, mitochondrial AAC is the one and only known ATP exporter that is evolutionarily distinct from other ATP/ADP translocases present in plastids and in intracellular pathogens that import ATP **(Amiri et al., 2003; Lin et al., 2025) (Figure 1)**.

**Figure 1.**
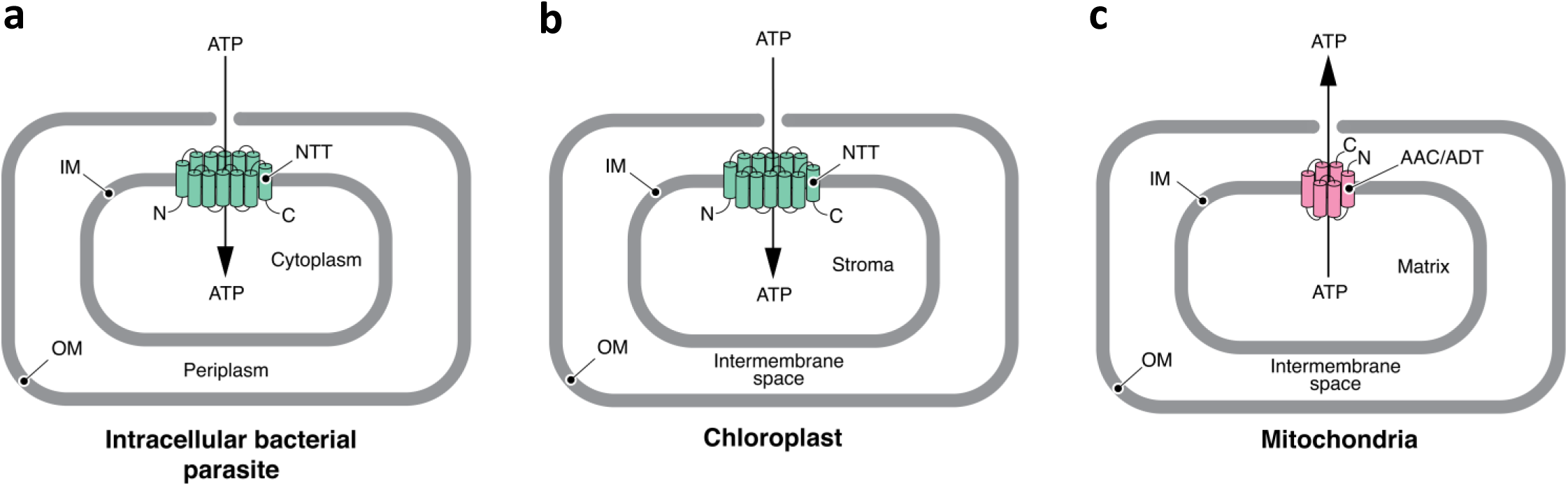
ATP transporter types. (**a**) Nucleotide transporter (NTT) present in intracellular bacterial parasite that imports ATP from host cell cytoplasm. Parasite NTT features all alpha 12 TM containing MFS general substrate transporter fold (SCOP id: 2000248). (**b**) ATP/ADP translocator 1 (NTT1) present in plastid inner membrane responsible for importing ATP from plant cell cytoplasm. Plastid NTTs feature all alpha 12 TM containing MFS general substrate transporter fold (SCOP id: 2000248). (**c**) Mitochondrial ATP/ADP translocator (ADT/ANT/AAC) that resides in mitochondrial and mitochondria derived organelles that exports ATP to the eukaryotic cell cytoplasm. AACs feature all alpha 6 TM bundle Mitochondrial carrier fold (SCOP id: 2000266).

Mitochondrial AACs belong to the nuclear encoded six transmembrane (6TM) Solute Carrier Family 25 (SLC25), also known as mitochondrial carrier family (MCF) **(Nury et al., 2006)**. SLC25 family is the largest mitochondrial carrier family constituted of 53 paralogues in *Homo sapiens*, 35 in *Saccharomyces cerevisiae*, 60 in *Arabidopsis thaliana* and 35 in *Andalucia godoyi* **(Palmieri et al., 2011)**. Metabolites transported by SLC25 across the impermeable mitochondrial inner membrane support myriad of eukaryotic processes, such as: oxidative phosphorylation, amino acid metabolism, apoptosis, iron-sulphur cluster synthesis, haem synthesis, heat production, mitochondrial fusion/fission dynamics, etc. **(Kunji et al., 2020; Ruprecht and Kunji, 2020; Kunji et al., 2025; Khan et al., 2025)**. AAC is the most extensively studied member of SLC25 family and for which the molecular mechanism of solute transport has been elucidated based on atomic structures of bovine and fungal AACs **(Pebay-Peyroula et al., 2003; Ruprecht et al., 2019, 2014)**. SLC25 transporters are conserved in all the extant branches of eukaryotes and were present in the last eukaryotic common ancestor (LECA) **(Kurland and Andersson, 2000)**. However, the evolutionary origin of the SLC25 mitochondrial carrier family, a crucial step in the emergence of mitochondria, remains an enigma owing to the lack of any trace of prokaryotic ancestry and therefore it is categorised as orphan or eukaryotic innovation **(Karlberg et al., 2000)**.

Here, we harness protein tertiary structure search to screen for putative remote homolog of AAC in the recently generated AlphaFold-predicted structure database of proteome of archaeal and bacterial species. This approach leverages the principle that protein tertiary structure is greater conserved over evolutionary distance compared to protein sequence. Our protein structure-guided screen identified bacterial proteins with significant structural similarity with AACs, which can be potential remote homologs of AACs in prokaryotes. Notably, the bacterial proteins tertiary structure is related to the structure of AACs through circular permutation of one transmembrane helix at sequence level. Furthermore, our thorough sequence analysis revealed the conservation of the hallmark SLC25 sequence motif in one among the structure search hits of AACs in bacteria, CysZ — a conserved bacterial inner membrane residing sulfate transporter protein. Thus, our comprehensive analysis identifies a bacterial homologue of mitochondrial AACs on the basis of structural similarity, functional similarity and conservation of a hallmark sequence-motif. Overall, identification of a bacterial ancestry establishes endosymbiotic roots for the emergence of the mitochondrial ATP exporter – a foundational step in the evolution of mitochondria and the onset of eukaryotic cell complexity.

## Results

### Rooted phylogeny resolves AACs as the founder member of SLC25 carrier family

Insights into the sequential order in which the members of SLC25 carrier family emerged is crucial for understanding the evolutionary emergence of AAC during the origin of mitochondria. Inferring the root of the phylogenetic tree of SLC25 family protein sequences may resolve the order in which AACs emerged relative to the rest of SLC25 family members. For phylogenetic analysis of SLC25 family, we took advantage of the SLC25 sequences of *Andalucia godoyi* **(Gray et al., 2020)**, a member of the eukaryotic order Jakobida under the supergroup Excavata, which is considered to be the most ancient branch of eukaryotes that is phylogenetically closest to LECA and harbours most bacterial-like mitochondria **(Lang et al., 1997; Burger et al., 2013; Williamson et al., 2025)**. We also employed SLC25 sequences from *Saccharomyces cerevisiae* and *Paramecium tetraurelia*, which belong to the eukaryotic supergroups Opisthokonta and SAR, respectively. The above three organisms from which we employ the SLC25 sequences to infer the phylogenetic tree are representative of three distinct supergroups of eukaryotes that have diverged at the stage of LECA **(Figure 2a) (Burki et al., 2020; Williamson et al., 2025)**. Next, we inferred phylogenetic tree of the SLC25 sequences using both

**Figure 2.**
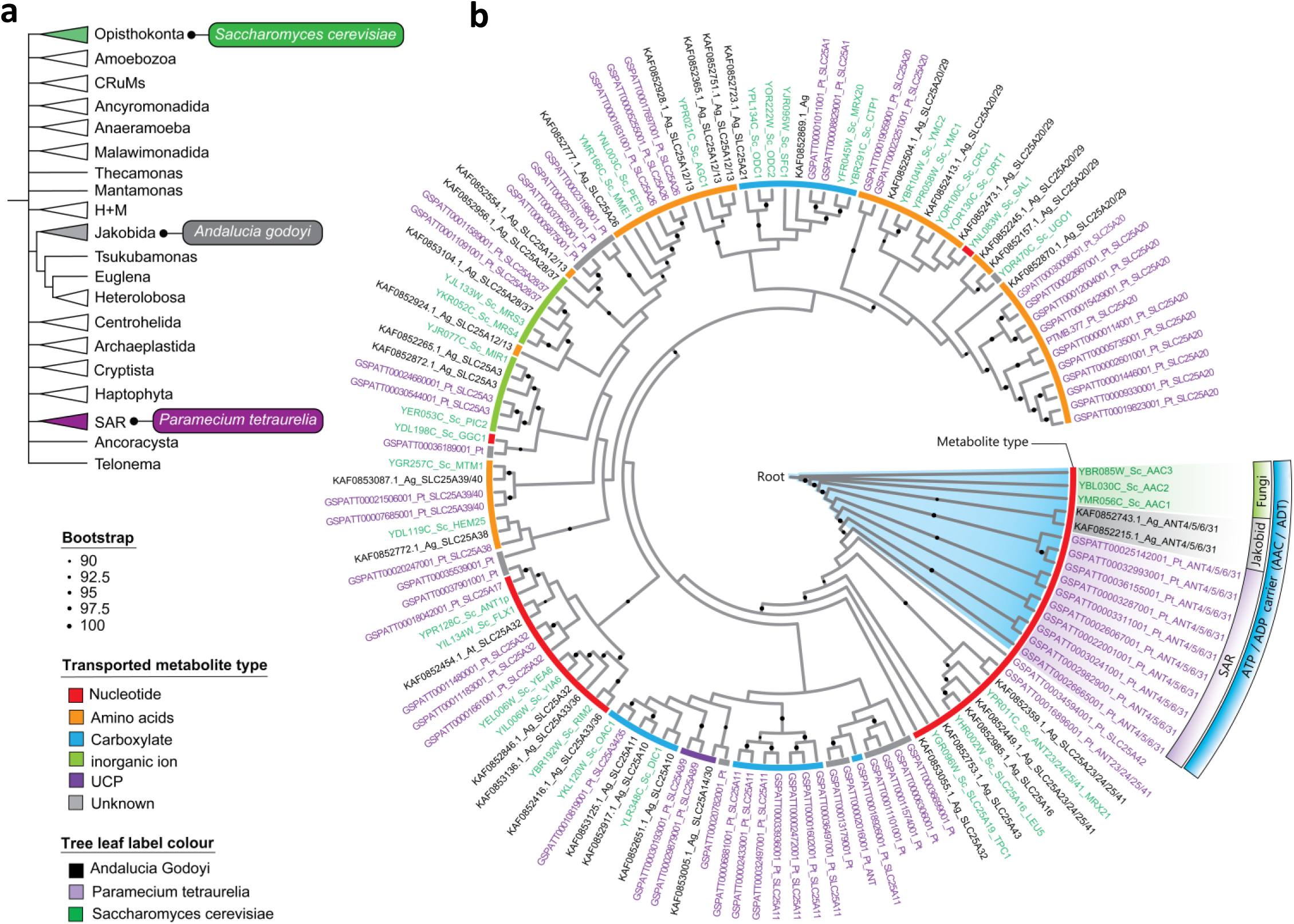
Mitochondrial ATP-ADP carriers (AACs) are the pioneer members of SLC25 family. **(a)** Tree showing all the Eukaryotic supergroups and the phylogenetic position of *A. godoyi*, *S. cerevisiae* and *P. tetraurelia* in three evolutionarily distinct eukaryotic supergroups that have diverged at the stage of LECA. **(b)** Maximum likelihood phylogenetic tree of SLC25 carrier family members from *A. godoyi*, *S. cerevisiae* and *P. tetraurelia*. Branch support of bootstrap value ≥ 90 are indicated by black circles.

Maximum likelihood (ML) and Bayesian methods **(Huelsenbeck and Ronquist, 2001; Nguyen et al., 2015)**. The ML phylogenetic tree of SLC25 carriers from *Andalucia godoyi*, *Paramecium tetraurelia* and *Saccharomyces cerevisiae* had all the AACs from the three organisms grouped together and also positioned at the root of the phylogenetic tree **(Figure 2b)**. Grouping is observed also for the remaining SLC25 transporters from the three organisms that transport similar metabolites. The Bayesian phylogenetic tree of the same sequences also exhibits a similar tree topology where the AACs are positioned at the root of the tree **(Supplementary Figure 1)**. Together, our comprehensive rooted phylogenetic analysis of SLC25 sequences from organisms belonging to three different eukaryotic supergroups that diverged at the stage of LECA resolves AACs as the evolutionarily founder member of the SLC25 carrier family.

### Structure-guided screen detects putative remote homologue of AAC in archaea and bacteria

Comparative genomics studies have established that the mitochondrial AACs is part of a group of proteins that are categorised as eukaryotic innovations due to lack of any trace of prokaryotic ancestry (**Karlberg et al., 2000**). Protein tertiary structure is significantly more conserved over evolutionary distance than sequence **(Holm and Sander, 1996; Illergård et al., 2009)**. Protein tertiary structure search therefore affords more confidence in detection of remote homologs well beyond the limits of sequence-based methods **(Murzin et al., 1995; Orengo et al., 1997; Cheng et al., 2014)**. Recent advancement in *ab initio* protein structure prediction programs like AlphaFold2 has enabled the generation of high quality AlphaFold structure database (AF-DB) of the entire known protein sequence space **(Baek et al., 2021; Jumper et al., 2021; Varadi et al., 2022)**. The predicted high quality structures in AF-DB allows for structure search-based screening of remote homologs in the *Foldome* of a target organism using programs like DALI AF-DB Search and Foldseek Search **(Holm, 2022; van Kempen et al., 2024)**. We set out to perform structure search of AACs using X-ray crystal structure of *Saccharomyces cerevisiae* AAC (*Sc*AAC) **(Pebay-Peyroula et al., 2003)** and high confidence AlphaFold3 **(Abramson et al., 2024)** predicted structure of *Andalucia godoyi* AAC (*Ag*AAC), in the predicted structures of the whole proteome of archaeal and bacterial species in the AF-DB using DALI AF-DB Search and Foldseek Search **(Figure 3a,b,c)**. Both *Ag*AAC and *Sc*AAC structures were searched in predicted structures of the entire proteome of bacterial species *E. coli* (Gram-negative), *S. aureus* (Gram-positive) and archaeal species through DALI AF-DB search. AAC structures were also searched across all archaeal and bacterial proteome structures in AF-DB through Foldseek Search. The structure search hits from the DALI AF-DB and Foldseek Search, having reviewed UniProtKB entry **(The UniProt Consortium, 2025)**, were filtered according to the degree of structural similarity to AAC based on DALI Z-score and TM-score cut-off criteria **(Holm and Sander, 1993; Zhang and Skolnick, 2005)**, and presence of overall 6TM topology based on DeepTMHMM **(Hallgren et al., 2022)** and manual curation **(Figure 3c)**. We retrieved five bacterial proteins and one archaeal protein that showed significant structural similarity to AACs, those that have DALI Z-score of more than 3 and TM-score of more than 0.4, and also features a 6TM fold topology like AACs **(Figure 3d,e,f and Supplementary Figure 2)**. Next, we performed sequence search to check the presence of each archaeal and bacterial hits across archaeal and bacterial phyla respectively **(Figure 4a)**. Only bacterial proteins YihY and CysZ were widely conserved across bacterial phyla—including Alphaproteobacteria—which asserts their likelihood of being the evolutionary ancestor of mitochondrial AACs; whereas, the rest of the archaeal and bacterial proteins were filtered out owing to their very confined distribution across selective prokaryotic phyla **(Figure 4b,c,d,e)**. Overall, our pipeline, comprising of protein tertiary structure search coupled with extensive curation based on phylogenetic conservation and transmembrane topology, identified bacterial proteins YihY and CysZ as potential remote bacterial homologs of AACs.

**Figure 3.**
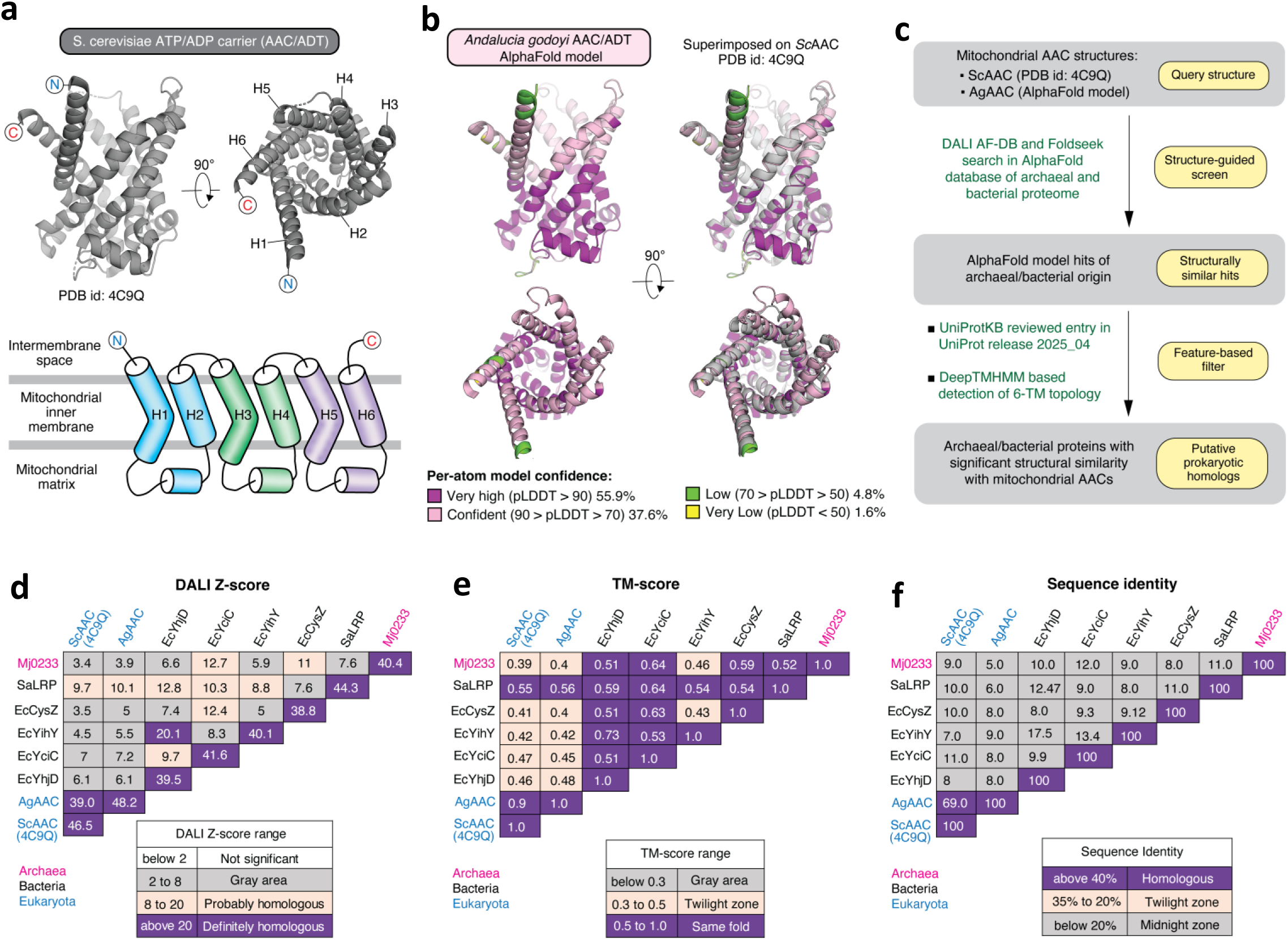
Structure guided screen of AACs in AlphaFold structure database of prokaryote proteome. **(a)** X-ray crystal structure of *Saccharomyces cerevisiae* AAC and its secondary structure topology. **(b)** AlphaFold predicted structure model of *Andalucia godoyi* AAC. **(c)** Schematic diagram of the pipeline employed in this study leveraging protein 3D-structrure search for identification of potential remote homologue of AACs in archaea and bacteria. **(d)** DALI Z-score, **(e)** TM-score, **(f)** sequence identity of archaeal and bacterial structure search hits of AACs.

**Figure 4.**
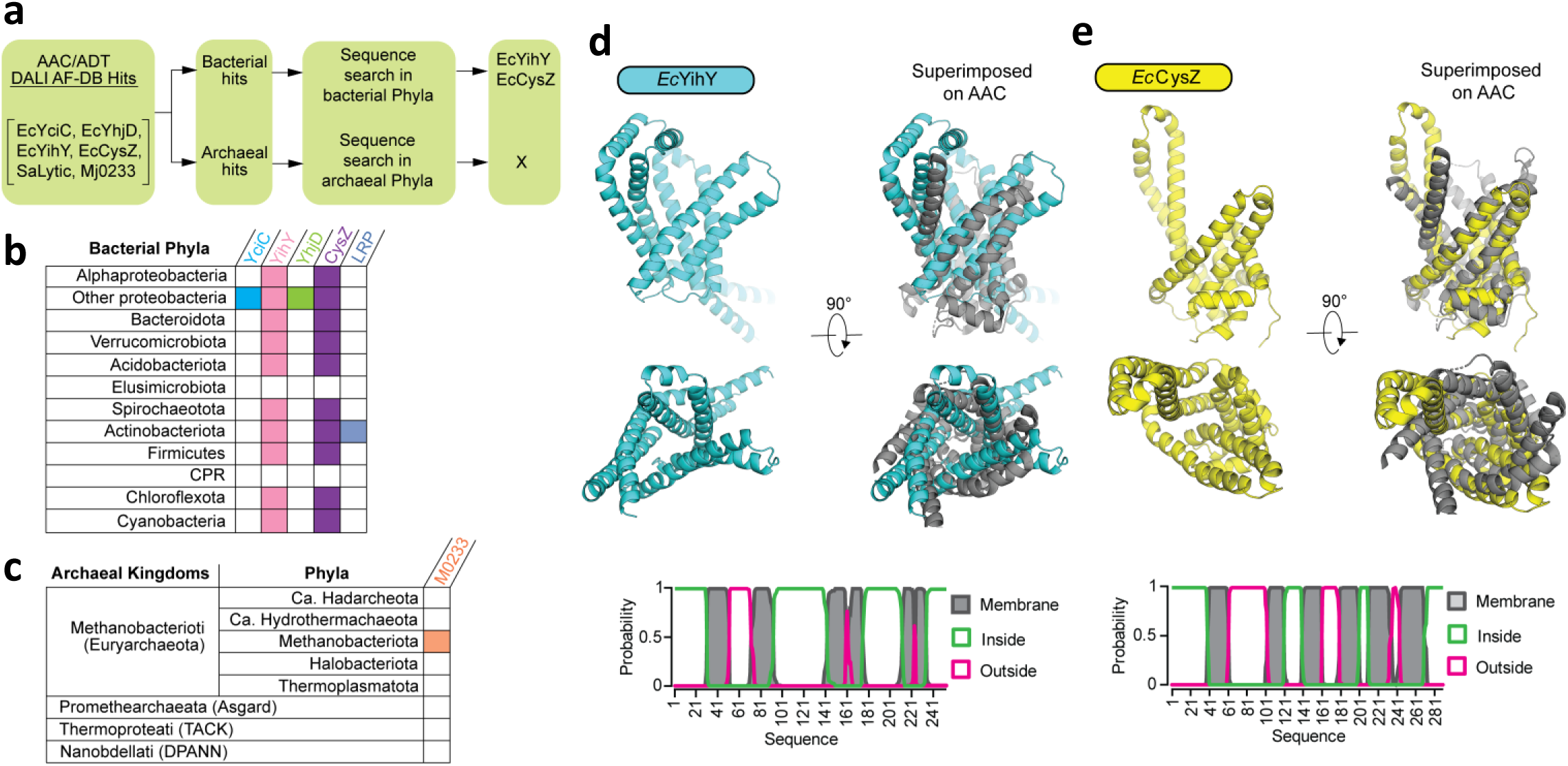
Putative homologue of AAC in prokaryotes. **(a)** Schematic flowchart depicting the approach for selection of the candidate homologue of AACs in prokaryote structural hits based on presence or absence of the structural hits in the bacterial and archaeal phyla. **(b)** Gene conservation of bacterial structural search hits of AAC across bacterial phyla. **(c)** Gene conservation of archaeal hits of structural search of AAC in archaeal kingdoms and phyla. **(d)** AlphaFold predicted model of *E. coli* YihY and its predicted transmembrane topology. **(e)** AlphaFold predicted model of *E. coli* CysZ and its predicted transmembrane topology.

### YihY and CysZ are structurally related to AACs through circular permutation

Structure of bacterial proteins YihY and CysZ is significantly similar to AACs, quantitated based on DALI Z-score of ∼ 5 and TM-score of ∼ 0.4, which both indicate probable homology **(Figure 3d,e)**. Inspection of the structural superimposition of YihY/CysZ and AACs reveal that although there is correspondence in the spatial arrangement of six TM helices in the structures, the connectivity of the helices in the primary structure is different **(Figure 5a)**. AACs feature a six TM-helical bundle (H1-6) with both N- and C-terminals present toward the intermembrane space (IMS). YihY and CysZ share a common structural fold featuring six TM-helical bundle with both N- and C-terminals in the cytoplasmic side. Inspection of the structural superimposition of YihY/CysZ and AAC reveal that there is circular permutation of the N- and C-terminal involving one TM helix in the protein sequence. The TM helix of YihY/CysZ structurally corresponding to the H6 of AAC has transposed to the N-terminal in the protein sequence, resulting in the shift of their N- and C-terminal at the position corresponding to the loop M1 of AAC, albeit maintaining a similar overall spatial arrangement of the six TM helices **(Figure 5b)**. Next, we sought to explore the effects of circular permutation of sequence on overall tertiary structure of YihY/CysZ. We computationally circular permuted YihY and CysZ by transposing their 1^st^ TM helix (h1) in the sequence to the C-terminal akin to the topology of AACs and then leveraged AlphaFold for prediction of their tertiary structure model **(Figure 5c,d,e)**. The AlphaFold predicted structure model of circular permuted proteins, YihY^CP^ and CysZ^CP^, superimposed on the AlphaFold model of native proteins with an RMSD of 0.38 Å over 290 Cα atoms and 0.29 Å over 253 Cα atoms respectively, suggesting that the inherent tertiary structure of this group of proteins is unaffected by alteration at sequence level by circular permutation **(Figure 5g,h)**. Furthermore, structure of circular permuted YihY^CP^ and CysZ^CP^ is more similar to AACs as indicated by the uptick in structure similarity scores compared to their native forms i.e., DALI Z-score increased from ∼ 4 to ∼ 6 and TM-score increased from ∼ 0.4 to ∼ 0.5 **(Figure 5i,j)**. Overall, our analysis reveals that the six TM helix bundle structure of YihY/CysZ and AACs are related by circular permutation of one TM helix, suggesting circular permutation as a potential step in the evolutionary route for the emergence of AACs from bacterial proteins YihY/CysZ.

**Figure 5.**
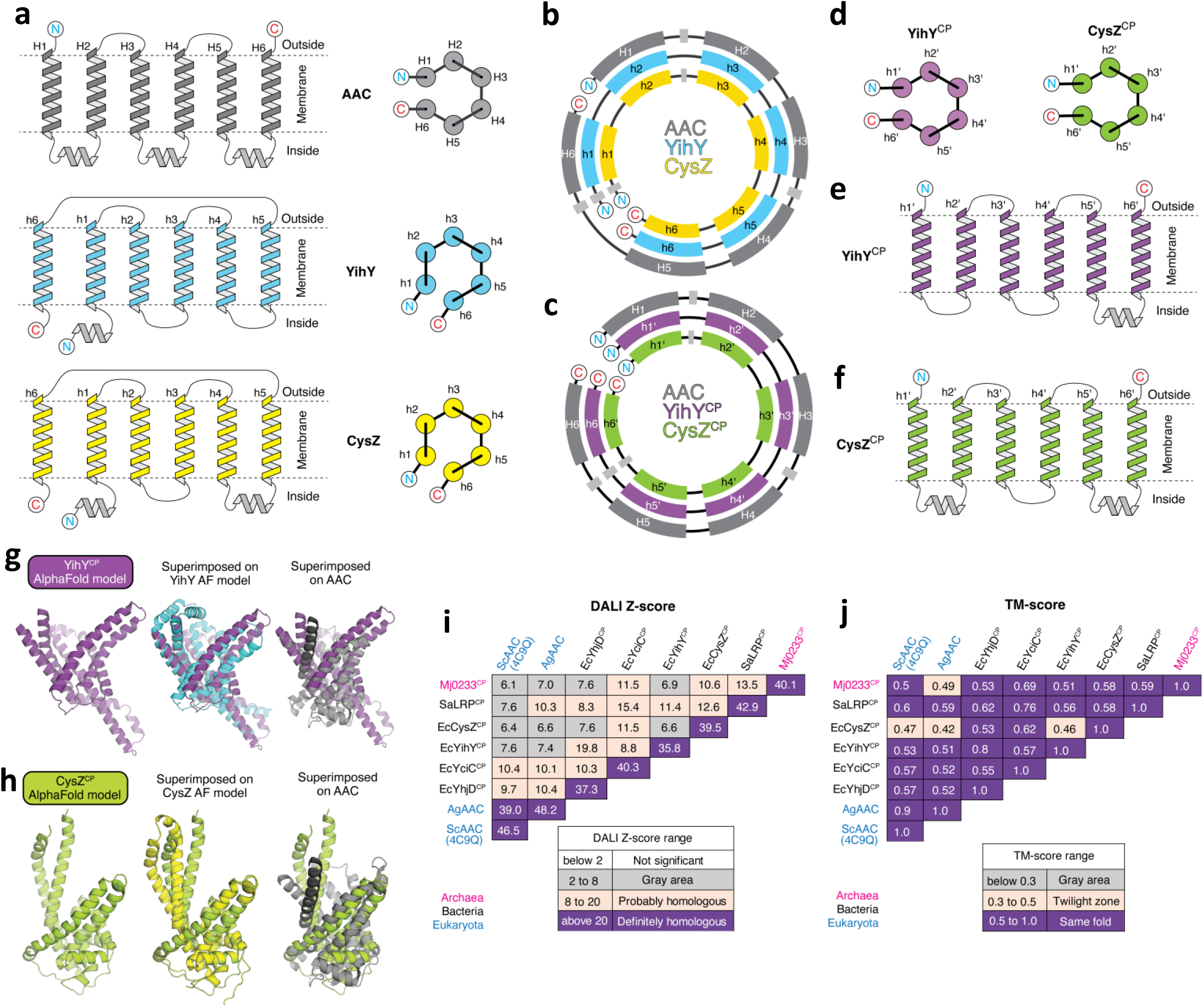
AAC is structurally related to prokaryotic proteins through circular permutation of a TM-helix. **(a)** Secondary structure topology and six TM-helix of AAC, YihY and CysZ (lateral and axial views). **(b)** Primary structure of AAC, YihY and CysZ showing circular permutation of one TM-helix. **(c)** Hypothetical circular permuted YihY^CP^ and CysZ^CP^ featuring similar secondary structure topology as to that of AAC. (**d**) Axial view of the six TM-helix topology of YihY^CP^ and CysZ^CP^. Secondary structure topology of (**e**) YihY^CP^ and (**f**) CysZ^CP^. **(g)** AlphaFold predicted structure model of YihY^CP^ and its superimposition on YihY and AAC. (**h**) AlphaFold predicted structure model of CysZ^CP^ and its superimposition on CysZ and AAC. **(i)** DALI Z-score and **(j)** TM-score for structural superimposition of AAC on AlphaFold predicted structure model of circular permuted bacterial and archaeal structure search hits.

### Presence of signature MCF motif in bacterial inner membrane sulfate transporter CysZ supports homology with AACs

Homology between two proteins means that both have descended from a common ancestor. Statistically significant sequence similarity between proteins alone is considered a definitive basis for assertion of homology. In absence of significant sequence similarity for highly diverged proteins, homology can be supported based on combination of structural resemblance, conservation of sequence motifs and functional similarity **(Grishin, 2001)**. In case of YihY/CysZ and AACs we show that they share significant structural similarity, which suggest for potential homology between them, but they are highly divergent at sequence level with sequence similarity score in the mid-night zone **(Figure 3d,e,f)**. In order to gain clarity regarding the hypothesised homology between YihY/CysZ and AACs, we examined their functional similarity and also checked for conservation of signature sequence motif. YihY (EcoCyc id: EG11851) is a conserved bacterial inner membrane protein of unknown function. CysZ (EcoCyc id: EG10003) is a conserved bacterial inner membrane protein, which has been shown to be a high specificity sulfate transporter involved in sulfate import inside cell for cysteine biosynthesis **(Assur Sanghai et al., 2018; Zhang et al., 2014; Britton et al., 1983)**. Thus, functionally CysZ is similar to AACs in that they both are membrane transporters of small metabolite.

Next, we sought to check for any motifs conserved between YihY/CysZ and AACs. All the members of SLC25 carrier family feature a conserved structure with six TM helices having a three-fold internal symmetry of a repeat containing two TM helices called as “mito_carr’ domain **(Figure 6a) (Pebay-Peyroula et al., 2003; Nury et al., 2006)**. The odd-numbered helix of each repeat features a highly conserved characteristic sequence motif – PX[D/E]XX[K/R]X[K/R], known as the MCF-motif, which forms inter-domain salt-bridges called as the matrix salt-bridge network. The matrix salt-bridge network is an integral part of the gates that has direct role in the alternate access mechanism of solute transport by SLC25 solute carrier family **(Ruprecht and Kunji, 2020)**. In case of YihY/CysZ, the six helical bundle can be assigned with repeat 2 and repeat 3 based on their mapping on the repeats of AACs in the structural overlap **(Figure 6b,c,d,e,f,g,h,i)**. In order to check for conservation of sequence motif between AACs and YihY/CysZ, we separated the sequences of the repeats of AACs and YihY/CysZ and then generated a structure-based multiple sequence alignment. Structure-based multiple sequence alignment of repeats revealed the conservation of the signature motif PX[D/E]XX[K/R]X[K/R] in the repeat 3 of CysZ **(Figure 6j,k and Supplementary Figure 3a)**.Thus, the bacterial sulfate transporter CysZ not only features a significantly similar structure like AACs, it also exhibits functional similarity and contains the signature MCF-motif, which altogether accentuate the support for their evolutionary relationship and thereby asserts homology between CysZ and AACs.

**Figure 6.**
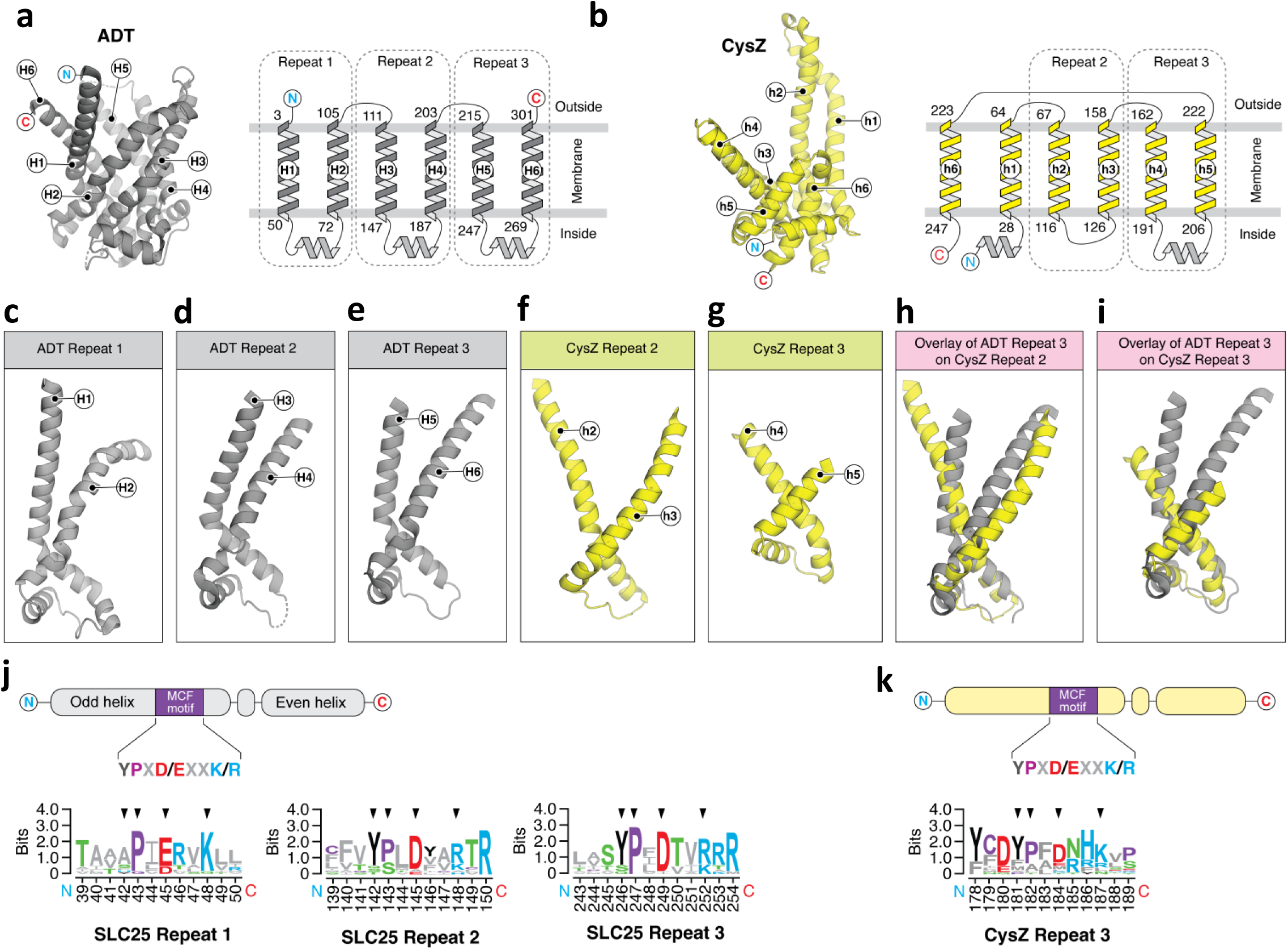
Conservation of MCF motif in CysZ. **(a)** Secondary structure topology of AAC and the three-fold internal symmetry of two TM-helix containing repeat. **(b)** Topology of CysZ and its two TM-helix repeats. Structure of **(c)** AAC repeat 1, (**d)** AAC repeat 2 **(e)** AAC repeat 3, **(f)** CysZ repeat 2, (**g**) CysZ repeat 3. **(h)** Structural overlay of AAC repeat 2 and CysZ repeat 2. **(i)** Structural overlay of AAC repeat 3 and CysZ repeat 3. **(j)** MCF motif in AAC repeat 1, 2 and 3. **(k)** MCF motif in CysZ repeat 3.

## Discussion

Evolutionary origin of mitochondrial ATP exporter, a foundational step in all the models of the evolution of mitochondria, remains unclear **(Gray, 2015)**. For decades, comparative genomics studies have categorised mitochondrial AACs and rest of the SLC25 family members as orphan or eukaryotic innovation owing to the lack of any trace of prokaryotic ancestry. Here, we first delineated the sequential order of evolutionary origin of the members of the SLC25 family through a comprehensive phylogenetic analysis and resolved AACs as the evolutionary founder member of SLC25 family. Then, we harnessed protein tertiary structure search of experimental as well as AlphaFold predicted structure of AACs in the AlphaFold predicted structure database of prokaryotic organisms, in tandem with thorough sequence analysis, to establish bacterial sulfate transporter CysZ as a potential evolutionary ancestor of mitochondrial AACs and the SLC25 carrier family.

AACs and all the other members of SLC25 carrier family features a three-fold pseudosymmetry of a two TM helical repeat **(Kuan and Saier, 1993)**. SLC25 members has been hypothesised to have evolved from multiple duplication of a two TM helical repeating unit, a process dubbed as repeat *triplication* **(Palmieri et al., 2011; Haferkamp and Schmitz-Esser, 2012)**. Due to the overall structural similarity and conservation of the signature MCF-motif, the repeat-3 of bacterial sulfate transporter CysZ makes for an ideal *arch*-repeat from which the AAC and the rest of the SLC25 carriers could have evolved through multiple duplication events.

Widely accepted syntrophy hypothesis and entangle-engulf-endogenize (E3) hypothesis for the emergence of proto-mitochondria are based on metabolic syntrophy which later transformed into energy exchange in form of ATP in mitochondria. The above metabolic syntrophy process involved sulfate as one of the metabolite exchanged between symbiont bacteria and the archaeal host **(Imachi et al., 2020; López-García and Moreira, 2020; Vosseberg et al., 2024)**. Our finding that the AAC emerged from the conserved bacterial sulfate transporter CysZ, strikingly, suggest that a transporter involved in the initial metabolic syntrophy that spurred the formation of proto-mitochondria was eventually evolved and re-purposed for ATP export in mitochondria at the stage of LECA **(Figure 7)**.

**Figure 7.**
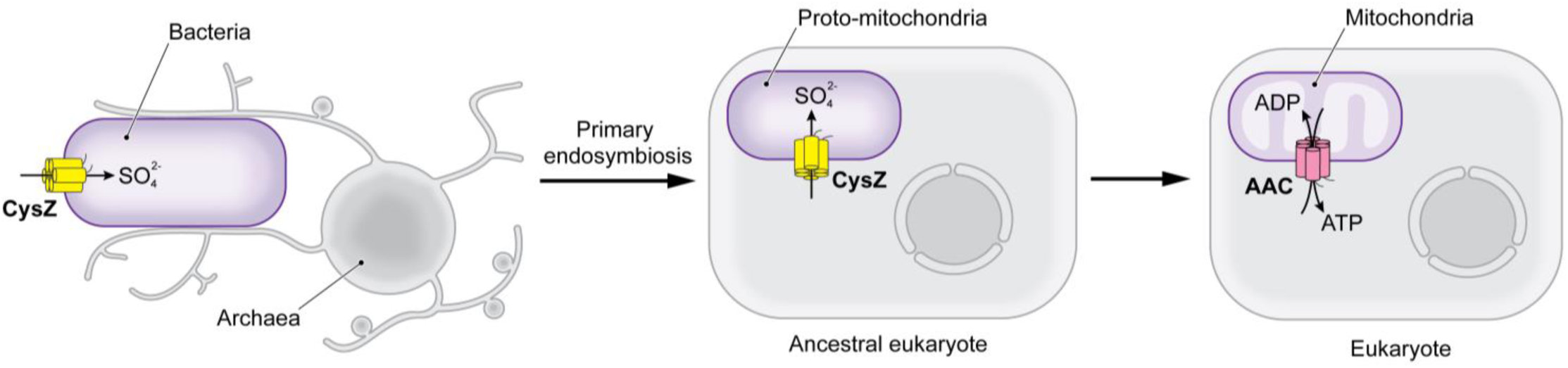
Model for evolution of mitochondrial AAC from bacterial CysZ. Schematic model depicting proposed origin of mitochondrial ATP exporter (AAC) from the bacterial sulfate transporter CysZ during the endosymbiotic origin of mitochondria in eukaryotic cells from bacterial endosymbiont. The bacterial sulfate transporter CysZ, involved in the initial metabolic syntrophy that spurred the formation of proto-mitochondria was eventually evolved and re-purposed for ATP export in mitochondria at the stage of LECA. Evolution of AAC from bacterial sulfate transporter CysZ is a foundational step of mitochondrial emergence, and therefore the onset of eukaryotic cell complexities.

About half of the ∼1000 mitochondrial targeted proteins lack any trace of archaeal or bacterial ancestry, which are categorised as orphan or eukaryotic innovation **(Kurland and Andersson, 2000; Gray, 2015)**. Various evolutionary mechanisms have been implicated for the evolution of this profound eukaryotic genomic novelties **(Vosseberg et al., 2024)**. Recently, optimization of the macromolecules for attaining compatibility among the bacterial and archaeal derived components for successful integration of various processes during the emergence of mitochondria was implicated in the emergence of the mosaic mitochondrial proteome **(Gogoi et al., 2022; Kumar et al., 2022, 2023)**. We speculate that many of the novel eukaryotic innovations that lack any trace of prokaryotic ancestry could be a result of extreme degree of optimization of the prokaryotic proteins, in the evolutionary course of emergence of the eukaryotic proteins, that the sequence level similarity among them is eroded beyond the detection capabilities of the sequence search methods. Here, we have leveraged the recently generated database of high quality AlphaFold predicted structure of the entire sequence space to screen for remote prokaryotic homolog of AACs using protein tertiary structure search programs. The retrieved structurally similar prokaryotic proteins facilitated implementation of focused sequence analysis methods to unravel the conserved MCF motif that unequivocally established the evolutionary relationship and therefore the homology between AAC and bacterial protein CysZ. We envisage that such an approach can be expanded to deorphanise many more such remote homologs that are categorised as orphan/eukaryotic innovation by unravelling the distant evolutionary relationship with their prokaryotic ancestors. Such studies may further shine a light on the evolutionary mechanisms that underpins the emergence of the eukaryotic genomic novelties and illuminate the origins of eukaryotic cellular complexities.

## Methods

### Sequence and annotation retrieval

All the SLC25 sequences from *Paramecium tetraurelia* and *Saccharomyces cerevisiae* were retrieved from KEGG GENES database using KEGG BLAST search (https://www.genome.jp/tools/blast/) of S. cerevisiae AAC3 (Uniprot id: P18238). SLC25 sequences of *Andalucia godoyi* were retrieved by performing Domain Enhanced Lookup Time Accelerated BLAST (DELTA-BLAST) of *S. cerevisiae* AAC3 (Uniprot id: P18238) sequence. The annotations details for each sequence were retrieved by searching the corresponding accession code in either Uniprot or NCBI Gene database.

### Sequence alignment

Structure-based multiple sequence alignment of the SLC25 sequences was performed using *Expresso* program in T-coffee server (Di Tommaso et al., 2011) using S. cerevisiae AAC structure (PDB id: 4C9Q) as structural template. Structure-based multiple sequence alignment of repeats was created by first separating the sequences of repeats from SLC25 and CysZ and then were aligned in *Expresso* program in T-coffee server by using structure of repeat extracted from S. cerevisiae AAC structure (PDB id: 2C9Q). Alignments were curated in Jalview (Waterhouse et al., 2009).

### Phylogenetic tree reconstruction

The Maximum likelihood phylogenetic tree was reconstructed from the structure-based multiple sequence alignment using online program IQ-tree (http://iqtree.cibiv.univie.ac.at/) (Nguyen et al., 2015). The Bayesian phylogenetic tree was reconstructed in using online program MrBayes (https://www.phylogeny.fr/) (Huelsenbeck and Ronquist, 2001). The phylogenetic trees were visualized in iTol (Letunic and Bork, 2019).

Atomic coordinates of the structures of AAC were downloaded from RCSB protein data bank. Structural superimposition of the structures was carried out in Coot using SSM method. Structural superimposition of bacterial and archaeal structures on AACs was carried out using CEAlign in PyMol. High quality predicted AlphaFold structure of proteins were retrieved from AlphaFold Protein Structure Database (https://alphafold.ebi.ac.uk/) (Varadi et al., 2022). Prediction of structures were carried out in AlphaFold Server (https://alphafoldserver.com/) (Abramson et al., 2024).

### Structure Search

Protein structure search was carried out in DALI AF-DB search (http://ekhidna2.biocenter.helsinki.fi/dali/) (Holm, 2022), and Foldseek Search server (https://search.foldseek.cojotinm/search) (van Kempen et al., 2024).

## Acknowledgements

We thank all the members of the RS’s Laboratory at CCMB for the critical discussions on the work and for careful review of the manuscript.

JG thanks University Grants Commission, India for research fellowship. RSN thanks healthcare theme projects-Fundamental and Innovative CSIR in Science of Tomorrow (FIRST; MLP-0162) and Niche Creation Project (NCP; MLP-0138) of CSIR, India; J.C. Bose Fellowship of SERB, India.

## Author contributions

JG and RSN designed the study. JG carried out the analysis. JG and RSN conceived and supervised the study. JG and RSN wrote the manuscript and reviewed it.

## Data and materials availability

Atomic coordinates of the *Saccharomyces cerevisiae* AAC structure used in the study was obtained from RCSB PDB under the Entry id: 4C9Q. All data needed to evaluate the conclusions in the paper are present in the paper and/or the Supplementary Material.

